# PipeMaster: inferring population divergence and demographic history with approximate Bayesian computation and supervised machine-learning in R

**DOI:** 10.1101/2020.12.04.410670

**Authors:** Marcelo Gehara, Guilherme G. Mazzochinni, Frank Burbrink

## Abstract

Understanding population divergence involves testing diversification scenarios and estimating historical parameters, such as divergence time, population size and migration rate. There is, however, an immense space of possible highly parameterized scenarios that are difsficult or impossible to solve analytically. To overcome this problem researchers have used alternative simulation-based approaches, such as approximate Bayesian computation (ABC) and supervised machine learning (SML), to approximate posterior probabilities of hypotheses. In this study we demonstrate the utility of our newly developed R-package to simulate summary statistics to perform ABC and SML inferences. We compare the power of both ABC and SML methods and the influence of the number of loci in the accuracy of inferences; and we show three empirical examples: (i) the Muller’s termite frog genomic data from Southamerica; (ii) the cottonmouth and (iii) and the copperhead snakes sanger data from Northamerica. We found that SML is more efficient than ABC. It is generally more accurate and needs fewer simulations to perform an inference. We found support for a divergence model without migration, with a recent bottleneck for one of the populations of the southamerican frog. For the cottonmouth we found support for divergence with migration and recent expansion and for the copperhead we found support for a model of divergence with migration and recent bottleneck. Interestingly, by using an SML method it was possible to achieve high accuracy in model selection even when several models were compared in a single inference. We also found a higher accuracy when inferring parameters with SML.

## Introduction

The process of population divergence and speciation is finally being realized across many non-model organisms with the use of genetic data and advanced statistical models. Understanding population divergence involves testing diversification scenarios and estimating historical parameters, such as divergence time, historical demography and migration rate (Nielsen & Beaumont, 2009)□. Under simple diversification scenarios it is possible to use the coalescent model (Kingman, 1982) with the likelihood function and MCMC to infer model probabilities and associated historical parameters (Beerli & Palczewski, 2010; Bouckaert et al., 2014; Gronau, Hubisz, Gulko, Danko, & Siepel, 2011; Hey, 2010; Yang & Rannala, 2010). There is, however, an immense space of possible diversification scenarios where several hypotheses may translate into complex, highly parameterized models that are difficult or impossible to solve analytically (Fagundes et al., 2007; Mayr, 1942)□.

To overcome these limitations, researchers have used alternative approaches to approximate posterior probabilities or marginal likelihoods of population parameters by reducing data to summary statistics (Beichman, Huerta-Sanchez, & Lohmueller, 2018)□. These summary statistics can be used in approximate Bayesian computation (ABC) and Supervised Machine Learning (SML) to test hypotheses in a flexible likelihood-free context. ABC uses simulations generated from parameter values sampled from prior probabilities to infer posterior probabilities by applying a rejection algorithm that discards all simulations where the distance to the observed data falls above an arbitrary tolerance level (Beaumont, 2011; Csilléry, Blum, Gaggiotti, & François, 2010). Alternatively, simulated summary statistics can be used in SML as training data (Schrider & Kern, 2018)□. For the simulated data, the link between population parameters and summary statistics is known, so the algorithm can learn this connection and infer model probability and parameter values for observed summary statistics (Burbrink & Gehara, 2018; Sheehan & Song, 2016). To perform ABC and SML, end-users need to create custom scripts to sample parameters from prior distributions and pass them to a simulator. This requires integration of many different packages in various languages and the user’s ability to control this workflow sets the limit on the testable diversity of scenarios and hypotheses.

ABC and SML algorithms were already implemented in different packages of the R statistical platform (Csilléry, François, & Blum, 2012; Kuhn, 2008). However, there is currently no R-package to generate simulations for simulation-based model inference. To fill that gap we developed a new R-package, called *PipeMaster*, that can be used to build models, add prior distribution to model parameters, and simulate coalescent data from these prior distributions. *PipeMaster* can also calculate summary statistics for the empirical data to allow statistical comparison between observed and simulated data.

Here we demonstrate the utility of our newly developed package for three empirical examples and evaluate the power of ABC versus SML and the influence of the number of loci in the accuracy of model inferences. In the first example, we tested 10 hypotheses of divergence for the Muller’s termite frog, *Dermatonotus muelleri*, using newly generated data of 2177 loci of ultra-conserved elements (UCE). In the second and third examples, we tested six different hypotheses for two species complexes of North American vipers, the cottonmouth and the copperhead, using pre-existent multi-locus data (Burbrink & Guiher, 2015). We show that *PipeMaster* can be used with other R-packages to perform model and parameter inference in a single platform and to test complex diversification hypotheses to better understand the evolution of organisms.

## Material and Methods

The PipeMaster R-package is currently available on github (www.github.com/gehara/PipeMaster) and can be installed via the *install_github* function from the devtools R-package. Below we describe the main features of the package and exemplify its use for model and parameter inference using empirical data with Nexgen and Sanger dimensions.

### The interactive menu

*PipeMaster* has an interactive menu that allows the user to build models and set up parameter priors. In addition, the *main.menu* function can take a *ms* simulator string (see Hudson, 2002 for more information about ms) for model specification, which can be generated interactively with the PopPlanner application (see Ewing, Reiff, & Jensen, 2015 for more information about this application)□. Alternatively, the user can input a tree topology in newick format as a backbone of a diversification model, thus generating a simple isolation model with constant population size and divergence time parameters. This basic isolation model can be modified by adding ancestral population size changes and migration parameters, or by removing divergence parameters to simulate island models. The user can use the interactive menu to set conditions for parameter sampling (e.g. Ne1 > Ne2: effective population size of population 1 is larger than effective population size of population 2). In the current version, uniform and normal prior distributions are allowed. When the user exits the menu, the model can be saved as an R object. A previously generated model object can be used as a template for a different model setup, eliminating the need to start from the beginning when generating a nested model. Specific characteristics of the data regarding number of base pairs and samples per population per gene can be obtained using the *get.data.structure* function. This function reads the parameters of the observed data and replicates them in the model.

### The simulation functions work-flow

*PipeMaster* uses *ms* (Hudson, 2002) as an internal R function,□ or *ms*ABC (Pavlidis & Laurent, n.d.)□ as the essential source of simulation. The program *ms* simulates coalescent trees under the Wright-Fisher model, and places segregating sites on these trees under the infinite site model.

*PipeMaster* has three simulation functions for non-hierarchical models: i) *sim.ms.sumstat*, used to simulate summary statistics optimized for Sanger-scale data; ii) *sim.coaltrees*, to simulate coalescent trees; and iii) *sim.msABC.sumstat*, to simulate summary statistics using the simulator *ms*ABC (Pavlidis, Laurent, & Stephan, 2010) as an external program (**Figure 1a-c**). All functions take as input the model object generated by the *main.menu* function. They have the same basic work-flow and are used to sample parameter values from prior distributions, convert the values to coalescent scale, pass those values to a coalescent simulator, and write the output in a text file. In the case of *sim.ms.sumstat*, the simulated data is passed to PopGenome R-package (Pfeifer, Wittelsburger, Ramos-Onsins, & Lercher, 2014) for summary statistics calculation and the entire simulation process is performed without calling any external program.

**Figure 1:**
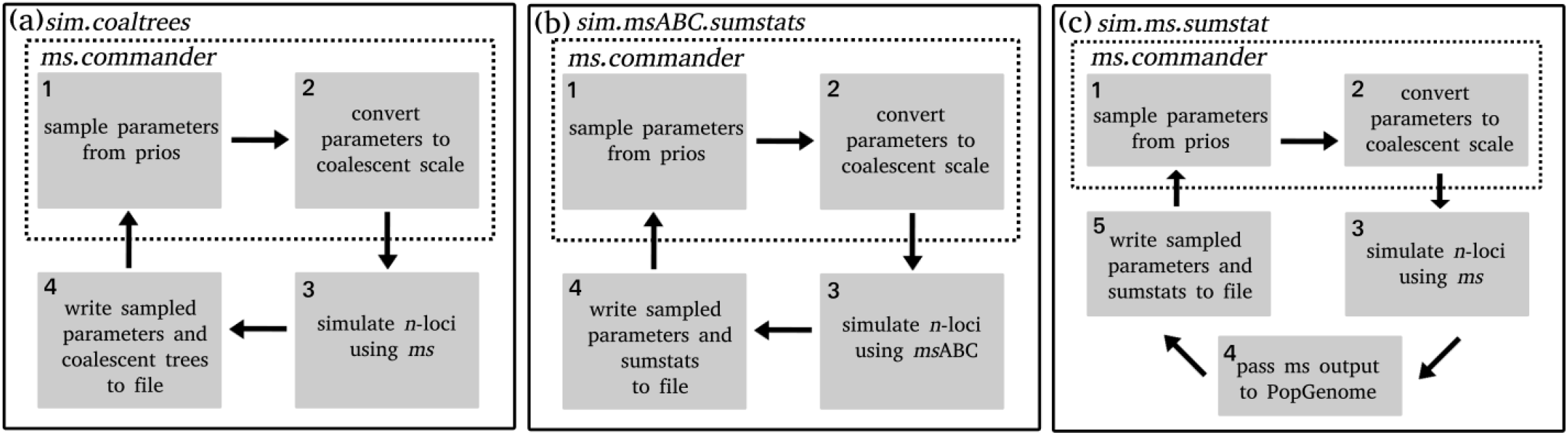
work-flow of the main simulation functions of PipeMaster and schematic representation of the simulated models in the toy example. (a) work-flow of the *sim.ms.sumstat* function; (b) schematic representation of the diversification models simulated in the toy example; (c) work-flow of the *sim.coaltrees* function; (d) work-flow of the *sim.msABC.sumstat* function.

### ABC and SML analyses

We implemented two different simulation-based inference methods in this study, approximate Bayesian computation (ABC) and supervised machine-learning (SML). In all empirical examples, before proceeding with the inference, we evaluated model-fit by running a PCA of simulated and observed data. For the ABC approach we used the *abc* R-package (Csilléry et al., 2012). We performed an abc rejection using the *postpr* function to calculate model probabilities by retaining 100 simulations with the closest distance from our observed data. To calculate accuracy in model selection we used *cv4postpr* with 100 pseudoreplicates per model and the same tolerance value. The final accuracy was calculated by dividing the total number of correct classifications by the total number of pseudoreplicates.

For SML we used the simulated data to train a neural network with one hidden layer to classify the data into different simulated scenarios using the *nnet* algorithm in *caret* R-package. We preprocessed the summary statistics by centering and scaling the data. We used 75% of the simulations as training data and the remaining 25% as testing data. To tune the parameters of the neural network, such as number of nodes and decay value, we performed 10 bootstraps with a maximum of 2,000 iterations in each learning replicate and retained the parameters yielding the highest accuracy. After training and testing, we used the neural network to classify our observed summary stats.

To estimate parameters we used the *abc* function of the abc R-package with the *neuralnet* regression method. Before proceeding with the estimation we simulated additional data for the best selected model totalizing 1,000,000 data sets for the Sanger examples and 100,000 for the UCE example. The *abc* function first performs a rejection step, reducing the dataset before neural network training. We evaluated tolerance and the accuracy of parameter estimates using the *cv4abc* function with 100 replicates and two different tolerance values for the Sanger examples (0.01, 0.001) and three values for the UCE example(0.1, 0.01, 0.001) .

We then calculated the correlation (*r*) between true and estimated parameters for each tolerance value. We selected the tolerance yielding the best correlations among parameters. All codes used in the ABC and SML are available on github as part of a tutorial for the package (github.com/gehara/PipeMaster).

### SML versus ABC and the influence of the number of loci in the accuracy of estimates

To evaluate the influence of the number of loci and compare the performance of ABC versus SML for estimating the true model, we ran a set of simulations experiments with four treatments that varied in total number of loci (10, 100, 1000, 2177). We used the case study of *Dermatonotus muelleri* below as an empirical basis for this experiment. Accordingly, simulation parameters, models, priors and summary statistics were the same as simulated for *D. muelleri*, while the different number of loci with their parameters (base pairs number of individuals per population) were obtained by sub-sampling 10, 100, or 1000 loci from the total dataset of 2177 loci generated for *D. muelleri*, plus a fourth treatment that contained the entire dataset. In each treatment we ran ABC and SML inferences for a group of pseudo-observed data (POD; i.e. test data in machine-learning jargon). We repeated these calculations three times, varying the total number of simulations per model (1,000, 10,000 and 100,000).

We also performed a simulation experiment based on the *D. muelleri* data to evaluate the accuracy of parameter estimates under different number of loci. In this case we estimated parameters for 100 POD under the IsBott2 model, which was the model with the highest probability for *D. muelleri* (see details below). We simulated a total of 100,000 data sets to use as reference data. To estimate parameters we used the *abc* function of the abc R-package with the neural network regression. We retained 1000 simulations after the rejection step and used these to train a neural network. We then calculated the correlation (*r*) between true value and estimated value for each parameter. An *r* closer to 1 would indicate a lower error in parameter estimates. We performed this calculation for the same treatments of 10, 100, 1000 and 2177 loci and we tested different retention or tolerance values.

### Application with UCE data - testing diversification hypotheses for muller’s termite frog

As an empirical example we generated a dataset of 2177 loci of ultra-conserved elements (UCE) for the neotropical frog, *Dermatonotus muelleri* (see details of molecular protocol in the Supplementary methods). This species is distributed along the dry diagonal of open formations which separates the Amazon from the Atlantic Forest. It is an explosive breeder, highly adapted to seasonal environments with pronounced periods of drought (Nomura, Rossa-Feres, & Langeani, 2009). A previous study using three loci (Oliveira et al., 2018)□ found that *D. muelleri* is composed by two deeply divergent populations, one distributed in the Caatinga and north of Cerrado, and a second one distributed in the southwest part of Cerrado (**Figure 2a**). Here we took a subsample of 88 individuals used in that study. After data assembly population assignment tests (see Supplementary methods) confirmed the existence of two spatially structured clusters (Oliveira et al., 2018)□.

**Figure 2:**
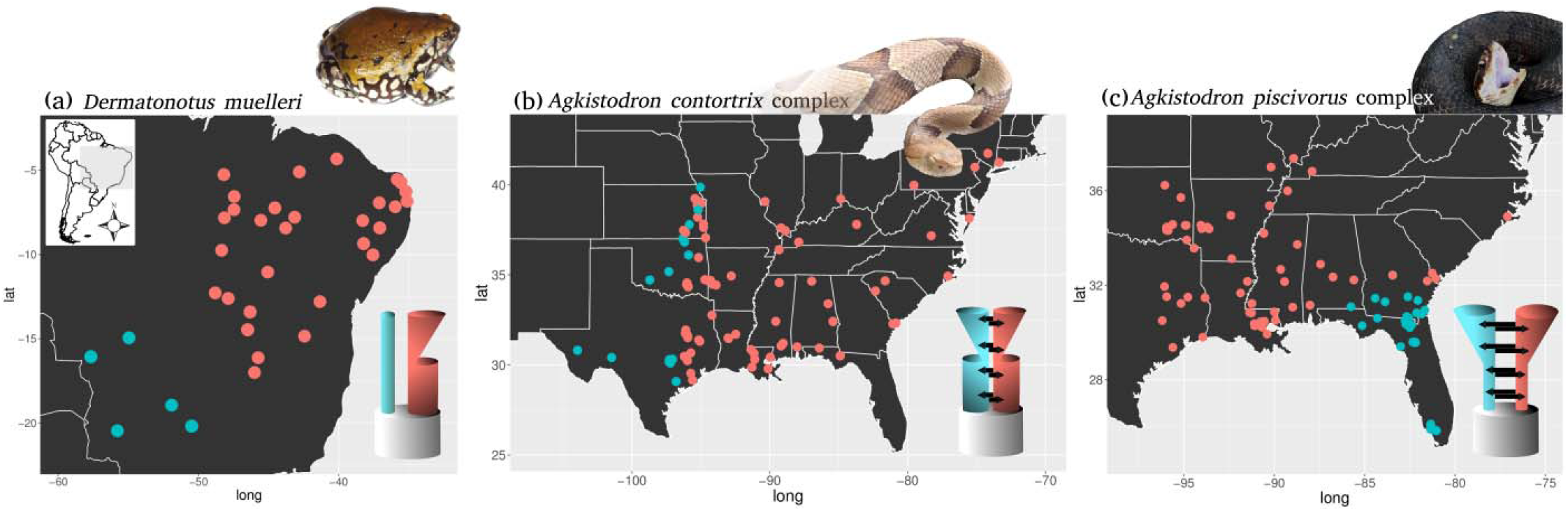
Distribution maps and best model for each data set analyzed in this study.

The geographic break separating these two populations falls in an area of high elevation, which may have isolated the populations. Also, Pleistocene climatic cycles are expected to have influenced the demographic history of at least the Northeast population (Gehara et al., 2017; Oliveira et al., 2018). Oliveira et al. (2018) found support for a model of diversification without migration and expansion only for the Northeastern population. To challenge these findings and test alternative diversification hypotheses for *D. muelleri*, we tested 10 two-population models: (i) a pure isolation scenario without migration and without demographic change (Is); (ii) an isolation with migration scenario without demographic change (IM); (iii) an isolation with recent expansion and no migration (IsExp); (iv) an isolation with migration and recent expansion (IMExp); (v) isolation with recent bottleneck and expansion (IsBott); (vi) isolation with migration, recent bottleneck and expansion (IMBott); (vii) isolation with recent expansion only for the Northeastern population (IsExp2); (viii) isolation with migration with expansion only for the Northeastern population (IMExp2); (ix) an isolation with bottleneck only for the Northeastern population (IsBott2); (x) an isolation with migration scenario with a bottleneck only for the Northeastern population (IMBott2) (**Figure 3**). Priors of population sizes and time of demographic events were retrieved from Oliveira et al. (2018) and can be found in the **Supplementary Table 1**.

**Figure 3:**
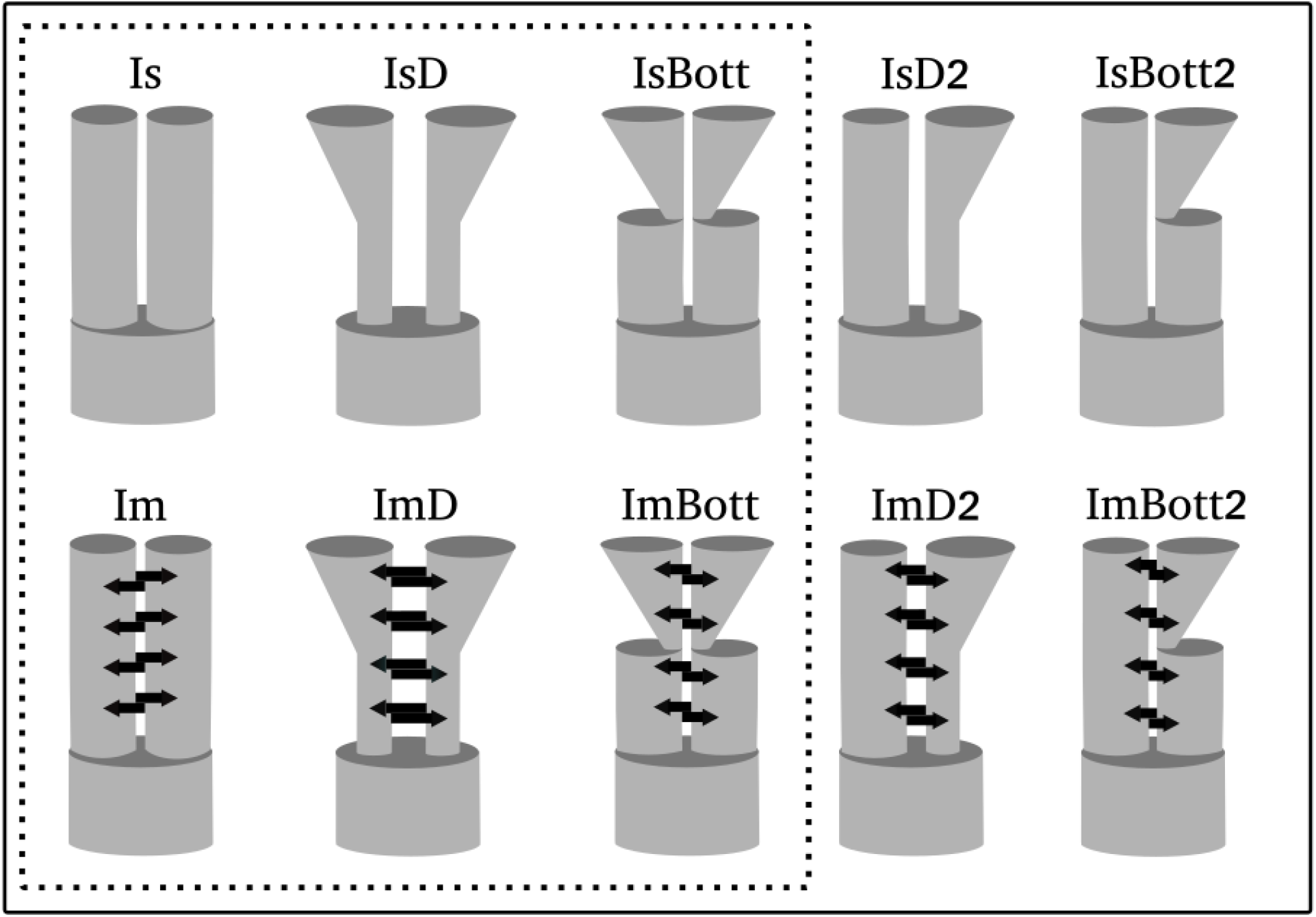
Schematic representation of the diversification models tested in the two *Agkistrodon* species complexes and *Dermatonotus muelleri*. Dotted line indicate the six models tested for *A. contortrix* and *A. piscivorus* complex. For *D. muelleri* we tested all 10 models. See **Supplementary Table 1** for a complete list of priors and parameters.

We simulated 100,000 data sets of 38 summary statistics (see **Supplementary Methods** and tutorial: github.com/gehara/PipeMaster) per model with *sim.msABC.sumstat* function. We used two independent approaches for model inference, ABC and SML described above.

### Application with Sanger data - testing diversification hypotheses for Copperhead and Cottonmouth pit vipers

We also performed a model selection for two species complexes of vipers widely distributed in Eastern North America: the *Agkistrodon contortrix* complex (Copperheads), and the *Agkistrodon piscivorus* complex (Cottonmouths). The dataset used contain one mitochondrial and five nuclear loci.

The *A. contortrix* species complex comprises two species, *A. contortrix* and *A. laticinctus*, which together cover a large portion of eastern and central United States. *Agkistrodon contortrix* is associated with deciduous hardwoods and pine forests and has a wider distribution in the Eastern and Midwestern US (**Figure 2b**). *Agkistrodon laticinctus* occurs in drier grassland environments in the central US to the Trans-Pecos habitats of west Texas. Diversification in this complex is likely ecological, since their contact zone falls in the transition from forested habitats to grasslands (Burbrink & Guiher, 2015). Both species currently occur in areas that were covered by ice sheet during the last glaciation and show genetic signs of population expansion in the Pleistocene (Guiher & Burbrink, 2008)□.

The *A. piscivorus* is also composed of two species. One of them, *A. conanti*, is mainly restricted to the Florida Peninsula. The other, *A. piscivorus*, is distributed north of the peninsula up to southern Illinois and Indiana in the north, Eastern Texas in the west, and coastal North Carolina in the east (**Figure 2c**). The contact zone of these two species in the Florida peninsula represents a common phylogeographic break for several other organisms (Burbrink, Fontanella, Alexander Pyron, Guiher, & Jimenez, 2008; Krysko, Nuñez, Lippi, Smith, & Granatosky, 2016; Mckelvy & Burbrink, 2017; Soltis, Morris, McLachlan, Manos, & Soltis, 2006)□ and the diversification of the complex was also likely influenced by the climatic cycles of the Quaternary (Guiher & Burbrink, 2008)□.

Taking these aspects into account, we tested for both species complexes, six diversification hypotheses (**Figure 3**). We generated the six models (a subset of the models simulated for the frog example above; see **Figure 3**) and simulated 100,000 datasets for each model using the *sim.ms.sumstat* function of PipeMaster R-package. We used wide uniform prior distributions according to Burbrink and Guiher (2015) (see parameter list and priors in the supplementary material). We used a set of 17 summary statistics (see **Supplementary Methods** and tutorial: github.com/gehara/PipeMaster)

For both species complexes and both methods used (ABC and SML) we compared the models hierarchically. (i) first we compared all the Isolation models with each corresponding version that included migration (e.g. IsD against IMD; IsBott against IMBott). (ii) Than we took the best models resulting from the first comparisons and conducted a second comparison to find the best model of all.

## Results

### SML versus ABC

The simulation experiment shows a higher error in model selection when using ABC relative to SML (**Figures 4 and 5**). The number of loci has a strong influence in the accuracy of model inferences. The dataset with 2177 loci had highest accuracy while the 10 locus dataset had the lowest. The number of simulations also influence accuracy with inferences performed with a reference dataset of 1,000 simulations per model having the lower true model probabilities (**Figure 4**), while the inferences performed with 100,000 simulations per model has the highest, particularly for ABC. For the SML inference both reference datasets of 10,000 and 100,000 simulations per model yielded nearly identical accuracies. The number of loci also has influence in parameter estimates. SML had higher precision when compared to ABC (**Figure 5**). The number of retained simulations, the tolerance value, influences ABC and SML in different ways. For ABC retaining a low number of simulations yielded higher *R*. For SML retaining more simulations result in better algorithm training.

**Figure 4:**
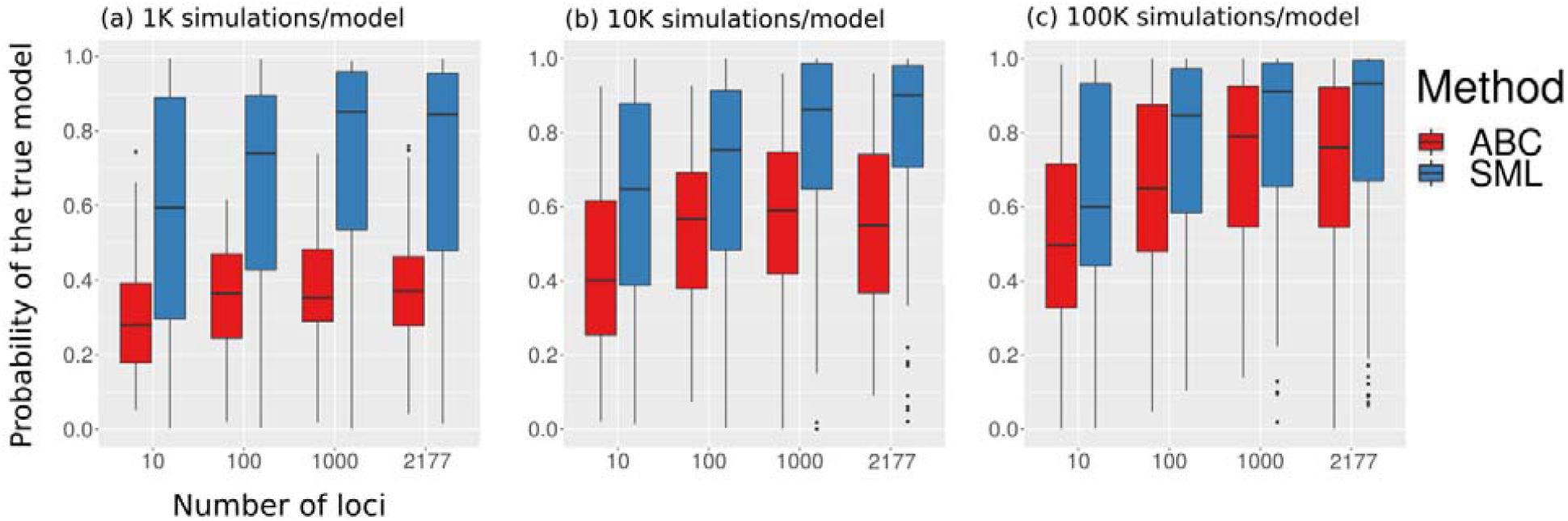
Results of the simulation experiment to compare the accuracy of ABC and SML for model inference in different conditions. The y-axis represents the probability of the true model, the x-axis represent different data dimensions. Each box plot represent probabilities of the true model for 100 pseudo observed data, 10 per model. For the ABC analysis, 100 simulations are retained in the rejection step, for the SML all simulations are used for algorithm training. (a) estimates performed with 1K simulations per model totalizing 10,000 simulations in the reference table. (b) estimates performed with 10K simulations per model totalizing 100,000 simulations in the reference table. (c) estimates performed with 100K simulations per model totalizing 1,000,000 simulations in the reference table.

**Figure 5:**
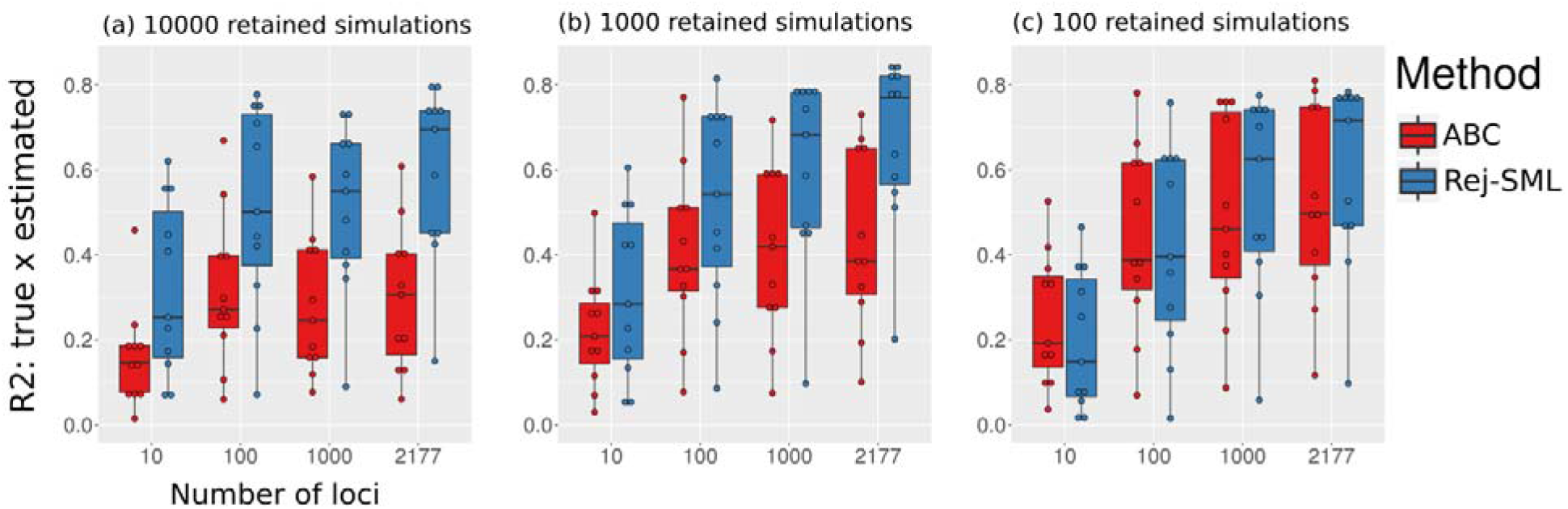
Results of the simulation experiment to evaluate the influence of number of loci and tolerance values on parameter estimates of ABC and rejection with SML. The y-axis represents the correlation between estimated and true values for 100 pesudo-observed data for the 11 parameters of the model. (a) estimates are performed by retaining 10,000 closest simulations. (b) estimates are performed by retaining 1,000 closest simulations. (c) estimates are performed by retaining 100 closest simulations.

### Diversification of muller’s termite frog

Simulations presented a good fit to the data as shown by the PCA plots (**Supplementary Figure 1**). The trained neural network is able to differentiate and classify the 10 models with an accuracy of 0.879 while the ABC had an accuracy of 0.83. Using the SML approach the observed data was classified as the IsBott2 model with a probability higher than 0.99 (**Table 1**), where only the northeast population experienced a bottleneck with expansion. The ABC inference suggest a different model, IMexp, where the two populations expand after divergence. In this case, the probability of the model was considerably lower, 0.49. Because the accuracy of the SML is higher, we consider the IsBott2 as the best model.

**Table 1:** Model probabilities and accuracies calculated with ABC and SML for the comparison of 10 simulated models for the frog *Dermatonotus muelleri* 2177 UCE data. (see **Figure 3** for a schematic representation of the models).

| Model | Probability |  |
| --- | --- | --- |
|  | SML | ABC |
| IM | 0 | 0 |
| IMBott | 0 | 0.17 |
| IMBott2 | 0 | 0 |
| IMExp | 0 | **0.49** |
| IMExp2 | 0 | 0 |
| Is | 0 | 0 |
| IsBott | 0.0037 | 0.17 |
| **IsBott2** | **0.9963** | 0.13 |
| IsExp | 0 | 0 |
| IsExp2 | 0 | 0.04 |
| **Accuracy** | **0.8798** | **0.826** |

The divergence time can be estimated with high accuracy and suggest a split around 2.6 Ma between the two populations (**Table 2**). Estimated current sizes for population 1 suggest a very large population after expansion but accuracy of this estimate is low. Estimates for population 2 are more accurate (**Table 2**). The average estimated mutation rate was 2.2E-10/site/generation with an estimated standard deviation of 3.88E-10/site/generation.

**Table 2:**
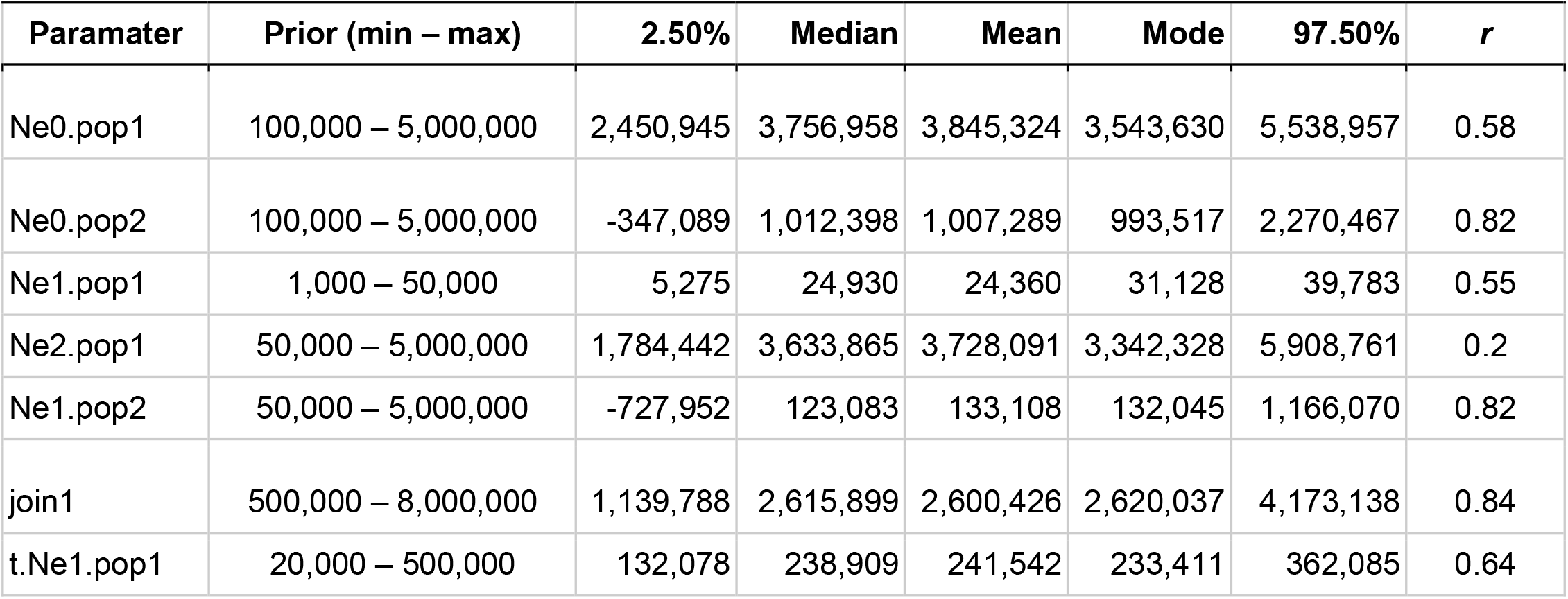
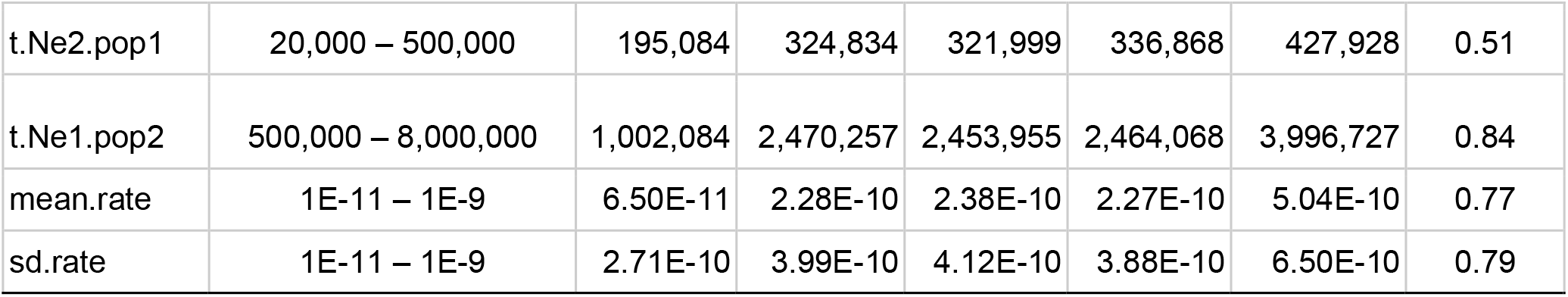
Parameter priors, posterior estimates and correlation (*r*) result calculated with the cross-validation experiment for the frog species (UCE data). See **Supplementary Table 1** for a complete list of priors and parameters.

### Diversification of Copperhead and Cottonmouth pit vipers

For both species complexes, the simulated models had a good fit to the data, as suggested by the PCA (**Supplementary Figure 2**). In the first comparisons (1, 2 and 3; see **Table 3**) for the *A. contortrix* complex, the accuracy varied from 0.79 – 0.85 for the SML and from 0.76 – 0.86 for the ABC. For comparison 1 (Is vs IM), ABC and SML showed conflicting results, with the pure isolation model, Is, having the highest probability for the ABC and the isolation with migration model, IM, having the highest probability in the SML. For comparisons 2 and 3, the two methods showed concordant results; models that included migration had higher probabilities than the correspondent models without migration (**Table 3**). The final comparison accuracies of ABC and SML were 0.78 and 0.79 respectively. Both methods converged in the same best model for the diversification of *A. contortrix* complex, IMBott (**Table 3**). For all comparisons, the SML showed higher probabilities for the selected model when compared to ABC (**Table 3**).

**Table 3:** Model probabilities estimated with ABC and SML with respective accuracies of estimates for the two snake species complex. Models were compared hierarchically, first comparisons 1, 2 and 3 were carried out independently. The final comparison included the best models of comparison 1, 2 and 3. Bold probabilities indicate the selected model for each comparison (see Figure 3 for a schematic representation of the models).

| *Agkistrodon piscivorus* |  |  |  |  |
| --- | --- | --- | --- | --- |
|  | 1 | 2 | 3 | Final |
|  | Is vs IM (Accuracy) | IsExp vs IMExp (Accuracy) | IsBott vs IMBott (Accuracy) | Best 1 vs Best 2 vs Best 3 (Accuracy) |
| SML | 0.11 / **0.89** (0.94) | 0.01 / **0.99** (0.94) | 0.04 / **0.96** (0.92) | 0.01 / **0.85** / 0.14 (0.79) |
| ABC | **0.78** / 0.23 (0.93) | 0.19 / **0.82** (0.91) | 0.49 / **0.51** (0.89) | 0.13 / **0.61** / 0.26 (0.87) |
| *Agkistrodon contortrix* |  |  |  |  |
|  | 1 | 2 | 3 | Final |
|  | Is vs IM (Accuracy) | IsExp vs IMExp (Accuracy) | IsBott vs IMBott (Accuracy) | Best 1 vs Best 2 vs Best 3 (Accuracy) |
| SML | 0.03 / **0.97** (0.79) | 0.00 / **1.00** (0.85) | 0.00 / **1.00** (0.79) | 0.12 / 0.01 / **0.87** (0.79) |
| ABC | **0.59** / 0.42 (0.76) | 0.08 / **0.92** (0.86) | 0.18 / **0.82** (0.77) | 0.33 / 0.01 / **0.66** (0.78) |

In the first comparisons (1, 2 and 3; see **Table 3**) for the *A. piscivorus* complex, the accuracy varied from 0.92 – 0.94 for the SML and from 0.89 – 0.93 for the ABC, and the best selected model were the same as the ones inferred for the *A. contortrix* complex (**Table 3**). In the final comparison, the accuracy of the ABC was higher than the SML, 0.87 and 0.79 respectively. However, both methods suggest high probabilities for the same model, the IMexp, which is an isolation with migration with expansion for both species (**Table 3**).

The cross-validation for the parameter estimates suggest low correlation between estimated and true values, particularly for *A. contortrix* (**Table 4**), suggesting high uncertainty in estimates. In general, the parameters that can be estimated with higher confidence are the current population sizes (**Table 4**).

**Table 4:**
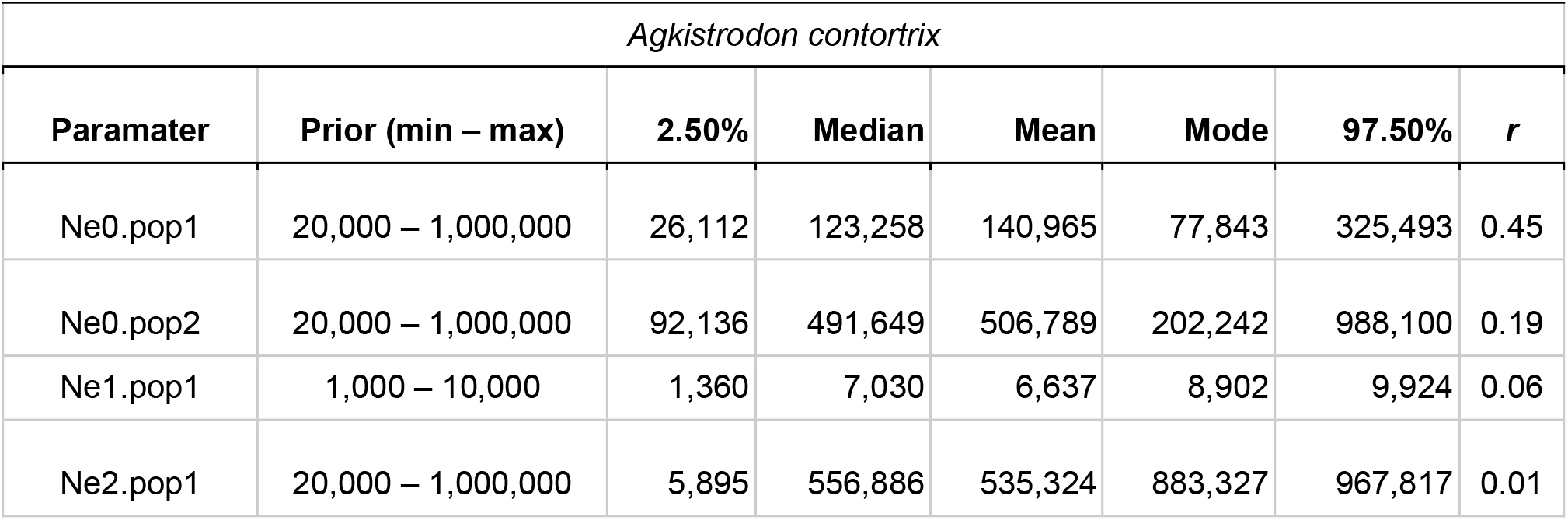
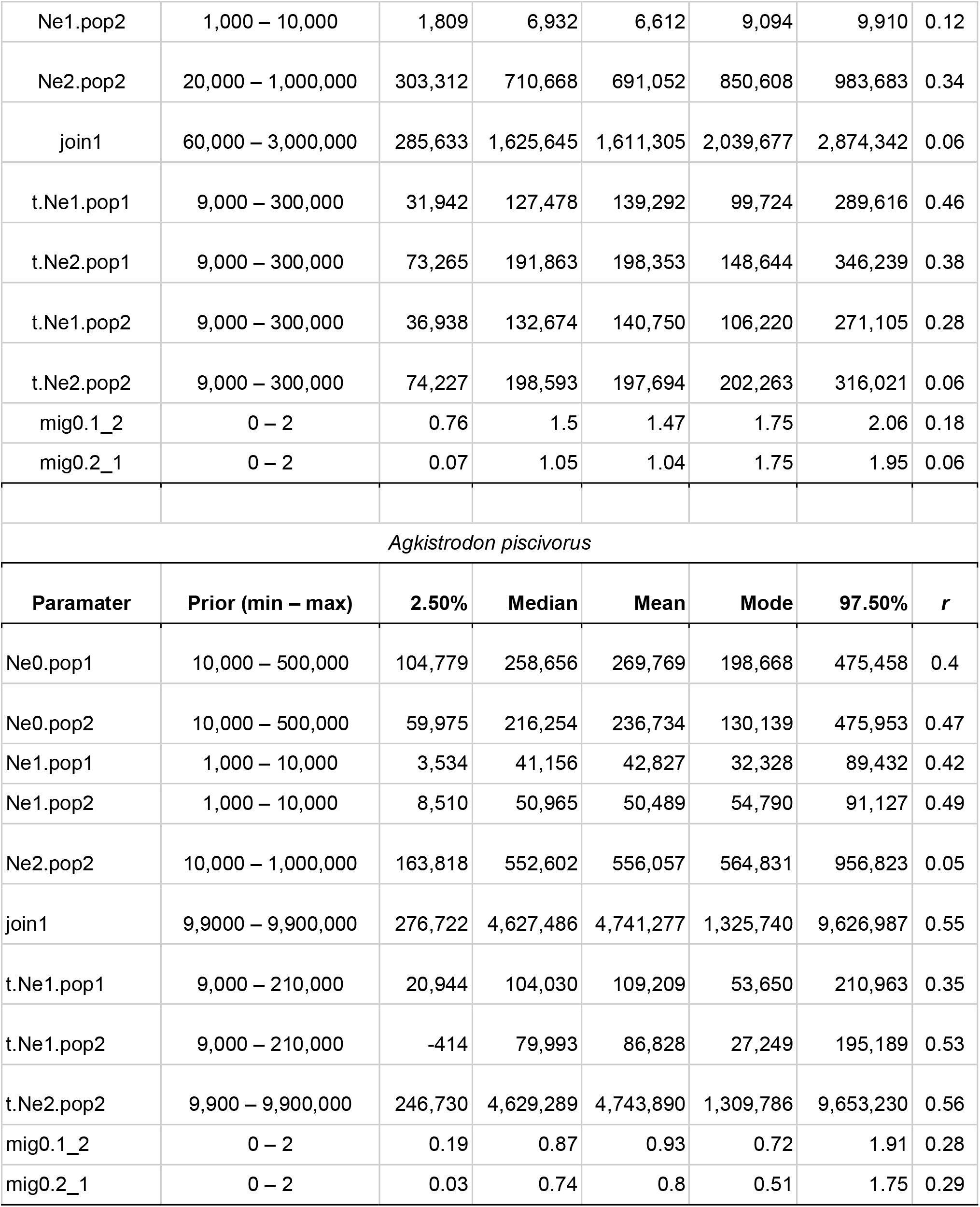
Parameter priors, posterior estimates and R result calculated with the cross-validation experiment for the two snake species complexes (Sanger data). See **Supplementary Table 1** for a complete list of priors and parameters.

| *Agkistrodon contortrix* |  |  |  |  |  |  |  |
| --- | --- | --- | --- | --- | --- | --- | --- |
| Paramater | Prior (min – max) | 2.50% | Median | Mean | Mode | 97.50% | r |
| Ne0.pop1 | 20,000 – 1,000,000 | 26,112 | 123,258 | 140,965 | 77,843 | 325,493 | 0.45 |
| Ne0.pop2 | 20,000 – 1,000,000 | 92,136 | 491,649 | 506,789 | 202,242 | 988,100 | 0.19 |
| Ne1.pop1 | 1,000 – 10,000 | 1,360 | 7,030 | 6,637 | 8,902 | 9,924 | 0.06 |
| Ne2.pop1 | 20,000 – 1,000,000 | 5,895 | 556,886 | 535,324 | 883,327 | 967,817 | 0.01 |
| Ne1.pop2 | 1,000 – 10,000 | 1,809 | 6,932 | 6,612 | 9,094 | 9,910 | 0.12 |
| Ne2.pop2 | 20,000 – 1,000,000 | 303,312 | 710,668 | 691,052 | 850,608 | 983,683 | 0.34 |
| join1 | 60,000 – 3,000,000 | 285,633 | 1,625,645 | 1,611,305 | 2,039,677 | 2,874,342 | 0.06 |
| t.Ne1.pop1 | 9,000 – 300,000 | 31,942 | 127,478 | 139,292 | 99,724 | 289,616 | 0.46 |
| t.Ne2.pop1 | 9,000 – 300,000 | 73,265 | 191,863 | 198,353 | 148,644 | 346,239 | 0.38 |
| t.Ne1.pop2 | 9,000 – 300,000 | 36,938 | 132,674 | 140,750 | 106,220 | 271,105 | 0.28 |
| t.Ne2.pop2 | 9,000 – 300,000 | 74,227 | 198,593 | 197,694 | 202,263 | 316,021 | 0.06 |
| mig0.1_2 | 0 – 2 | 0.76 | 1.5 | 1.47 | 1.75 | 2.06 | 0.18 |
| mig0.2_1 | 0 – 2 | 0.07 | 1.05 | 1.04 | 1.75 | 1.95 | 0.06 |
| *Agkistrodon piscivorus* |  |  |  |  |  |  |  |
| **Paramater** | **Prior (min – max)** | **2.50%** | **Median** | **Mean** | **Mode** | **97.50%** | ***r*** |
| Ne0.pop1 | 10,000 – 500,000 | 104,779 | 258,656 | 269,769 | 198,668 | 475,458 | 0.4 |
| Ne0.pop2 | 10,000 – 500,000 | 59,975 | 216,254 | 236,734 | 130,139 | 475,953 | 0.47 |
| Ne1.pop1 | 1,000 – 10,000 | 3,534 | 41,156 | 42,827 | 32,328 | 89,432 | 0.42 |
| Ne1.pop2 | 1,000 – 10,000 | 8,510 | 50,965 | 50,489 | 54,790 | 91,127 | 0.49 |
| Ne2.pop2 | 10,000 – 1,000,000 | 163,818 | 552,602 | 556,057 | 564,831 | 956,823 | 0.05 |
| join1 | 9,9000 – 9,900,000 | 276,722 | 4,627,486 | 4,741,277 | 1,325,740 | 9,626,987 | 0.55 |
| t.Ne1.pop1 | 9,000 – 210,000 | 20,944 | 104,030 | 109,209 | 53,650 | 210,963 | 0.35 |
| t.Ne1.pop2 | 9,000 – 210,000 | -414 | 79,993 | 86,828 | 27,249 | 195,189 | 0.53 |
| t.Ne2.pop2 | 9,900 – 9,900,000 | 246,730 | 4,629,289 | 4,743,890 | 1,309,786 | 9,653,230 | 0.56 |
| mig0.1_2 | 0 – 2 | 0.19 | 0.87 | 0.93 | 0.72 | 1.91 | 0.28 |
| mig0.2_1 | 0 – 2 | 0.03 | 0.74 | 0.8 | 0.51 | 1.75 | 0.29 |

## Discussion

Our simulation experiment showed that supervised machine-learning outperforms approximate Bayesian computation. This is particularly evident for datasets with genomic dimentions, which is the current standard of molecular studies for non-model organisms. We also show that much higher accuracies can be obtained with a SML as opposed to ABC, even when using just 100 loci and a considerably low number of simulations per model (10,000). Thus, because ABC requires a larger amount of simulations, it is more time consuming and less efficient when compared to SML.

Our simulation experiment also show that the model parameters can be estimated with higher accuracy with the increase in the number of loci. The SML approach also outperforms ABC for parameter estimates (**Figure 4**). Some parameters, like current effective population size and time of divergence, can be estimated with higher accuracy. However, ancestral population sizes are harder to estimate (**Figure 5**; **Table 2**). Interestingly, posterior distributions of the average and standard deviation of the mutation rate across all loci can be obtained with high confidence, allowing a more relaxed assumption when compared to using a fixed mutation rate for all loci.

### Diversification of muller’s termite frog

We found support for an isolation model with population contraction with expansion for the northeast population. This partially agrees with Oliveira et al. (2018), who found support for recent expansion without a contraction. Oliveira et al. (2018) analyzed only three loci while we analyzed more than 2,000, thus our data certainly contains more information about historical demography □(e.g. Gill et al., 2013).

The inference of a population contraction in the northeast population reinforces the idea of dynamic landscape changes in the northeast of Brazil along the Pleistocene. Currently, this area is predominantly covered by the Caatinga semiarid environment, but many studies suggest periods of increase in humidity in the last 1 Ma (Auler et al., 2004b; Cheng et al., 2013)□. Travertine deposits suggest a long period of increase in humidity from approximately 460 to 330 K years (Auler et al., 2004a)□ which remarkably agrees with our estimated time for the reduction in population size (mode: 337 Ky, CI: 195 – 437 Ky). These humid phases in the northeast of Brazil may have allowed long distance dispersals between Amazon and Atlantic forest fauna (Dal Vechio, Prates, Grazziotin, Zaher, & Rodrigues, 2018; Prates, Rivera, Rodrigues, & Carnaval, 2016). The reduction in population size is followed by a population expansion starting at around 230 K years (CI: 132 – 362 K years), in agreement of other studies that find synchronous population expansion Caatinga’s herpetofauna (Gehara et al., 2017)□.

The estimated divergence time at 2.6 Ma is considerably younger than previous estimates (∼4 Ma; see Oliveira et al., 2018). Our estimates places the divergence between the northeast and southwest populations in the Pliocene-Pleistocene transition, after the mid Pliocene warm period, when the average global temperature was 2 – 3^°^ C higher than today. This higher temperature may have allowed *D. muelleri*, a lowland species, to inhabit the highlands of the Brazilian plateau. With the temperature cooling the highland climate may have become unsuitable for the species and the Brazilian plateau became a vicariant barrier causing diversification.

### Diversification of Copperhead and Cottonmouth pit vipers

For both species complexes, we found support for demographic change and gene flow between species pairs. For the *A. contortrix* complex, we found support for a reduction in population size with subsequent expansion in the late Pleistocene. This species complex is currently found in areas that were covered by ice sheets during glaciations. Accordingly, the glaciation cycles would have restricted the distribution of the species to southern refugia, causing a population contraction (Burbrink et al., 2016; Marshall, James, & Clarke, 2002)□. In interglacial periods, the species would expand their range and their population sizes. It is also possible that the climatic cycles influenced their divergence, driving speciation by the isolation of populations in distinct refugia. Nevertheless, the presence of gene flow indicates that if isolation happened during glaciations, they were likely followed by periods of contact. Gene flow may also indicate the role of climatic gradients in diversification. *Agkistrodon contortrix* and *A. latiscinctus* occur in distinctly different niches (Burbrink & Guiher, 2015; Gloyd & Conant, 1990) and they likely present physiological adaptations to these different environments. Thus, hybrids may have lower fitness when compared to non-hybrids (Gow, Peichel, & Taylor, 2007)□. Future studies using thousands of loci will have the opportunity to test for selection across the climatic gradients, and may shed more light on the evolution of the *A. contortrix* species complex.

For the *A. piscivorus* complex, we found no support for a bottleneck during the Pleistocene. The most probable model suggests an isolation with gene flow and a recent population expansion. Both *A. piscivorus* and *A. conanti* are mostly distributed in areas free from broadscale effects of Pleistocene glaciation (Marshall et al., 2002). Accordingly, the supported model suggests a relatively more stable population size, with recent population expansion for both species. The contact zone between the species is in the northern area of the Florida Peninsula. This region was isolated from the continent when sea levels were higher, so it is likely that the diversification of the complex was influenced by sea level rise, which could have isolated *A. conanti* in a continental island formed by part of the landmass that today represents the Florida Peninsula (Hine, 2013; Krysko et al., 2016)□. In this scenario, gene flow between *A. conanti* and *A. piscivorus* was favored during glacial periods when sea levels were low, while isolation happened during interglacial periods while sea levels were high.

## Conclusion

We demonstrated the use of coalescent simulations generated by our newly developed R-package to infer the probability of complex diversification models in three different non-model organisms. In the three cases, we were able to test relatively complex demographic models with population size change, population structure and migration that are difficult, time consuming or impossible to implement using a full Bayesian or likelihood approaches. Interestingly, by using a SML method it was possible to achieve high accuracy in model selection even when several models were compared in a single inference (**Table 1**).

Machine-learning algorithms are becoming increasingly available to the general scientific community through R and Python applications, facilitating its use for an unprecedented number of cases in evolutionary biology and ecology. Here we demonstrated its use comparing it with a more traditional, ABC, for model inference in population genetics. Our results agree with the recent literature (Schrider & Kern, 2018; Sheehan & Song, 2016)□ supporting the power of SML in dealing with complex multi-dimensional problems such as the ones presented here.

## Acknowledgments

We thank Vinicius A. São Pedro, Eliana Oliveira and Adrian A. Garda for helping gathering the data; and Brian T. Smith, Gustavo Cabanne and Gregory Thom e Silva for trying beta versions of the package.

## Author Contributions

MG and FB conceived the ideas; MG, GGM and FB designed methodology; FB and MG collected the data; MG analyzed the data; MG and FB led the writing of the manuscript. All authors contributed critically to the drafts and gave final approval for publication.

## Data Availability

All codes used in the ABC and SML analyses are found in github.com/gehara/PipeMaster. The assembled UCE data is available in the Dryad (upon manuscript acceptance).

## References

Auler, A. S., Wang, X., Edwards, R. L., Cheng, H., Cristalli, P. S., Smart, P. L., & Richards, D. A. (2004a). Palaeoenvironments in semi-arid northeastern Brazil inferred from high precision mass spectrometric speleothem and travertine ages and the dynamics of South American rainforests. Speleogenesis and Evolution of Karst Aquifers, 2(2), 1–4.

Auler, A. S., Wang, X., Edwards, R. L., Cheng, H., Cristalli, P. S., Smart, P. L., & Richards, D. A. (2004b). Quaternary ecological and geomorphic changes associated with rainfall events in presently semi-arid northeastern Brazil. Journal of Quaternary Science, 19(7), 693–701.

Beaumont, M. A. (2011). Approximate Bayesian Computation in Evolution and Ecology. Annual Review of Ecology, Evolution, and Systematics, 41(1), 379–406.

Beerli, P., & Palczewski, M. (2010). Unified framework to evaluate panmixia and migration direction among multiple sampling locations. Genetics, 185(1), 313–326.

Beichman, A. C., Huerta-Sanchez, E., & Lohmueller, K. E. (2018). Using Genomic Data to Infer Historic Population Dynamics of Nonmodel Organisms. Annual Review of Ecology, Evolution, and Systematics, 49(1), annurev – ecolsys – 110617–062431.

Bouckaert, R., Heled, J., Kühnert, D., Vaughan, T., Wu, C.-H., Xie, D., … Drummond, A. J. (2014). BEAST 2: a software platform for Bayesian evolutionary analysis. PLoS Computational Biology, 10(4), e1003537.

Burbrink, F. T., Chan, Y. L., Myers, E. A., Ruane, S., Smith, B. T., Hickerson, M. J., & Sgro, C. (2016). Asynchronous demographic responses to Pleistocene climate change in Eastern Nearctic vertebrates. Ecology Letters, Vol. 19, pp. 1457–1467. doi: 10.1111/ele.12695

Burbrink, F. T., Fontanella, F., Alexander Pyron, R., Guiher, T. J., & Jimenez, C. (2008). Phylogeography across a continent: The evolutionary and demographic history of the North American racer (Serpentes: Colubridae: Coluber constrictor). Molecular Phylogenetics and Evolution, 47(1), 274–288.

Burbrink, F. T., & Gehara, M. (2018). The Biogeography of deep time reticulation. Systematic Biology, 67(5), 743–744.

Burbrink, F. T., & Guiher, T. J. (2015). Considering gene flow when using coalescent methods to delimit lineages of North American pitvipers of the genus Agkistrodon. Zoological Journal of the Linnean Society, 173(2), 505–526.

Cheng, H., Sinha, A., Cruz, F. W., Wang, X., Edwards, R. L., D’Horta, F. M., … Auler, A. S. (2013). Climate change patterns in Amazonia and biodiversity. Nature Communications, 4, 1411.

Csilléry, K., Blum, M. G. B., Gaggiotti, O. E., & François, O. (2010). Approximate Bayesian Computation (ABC) in practice. Trends in Ecology & Evolution, 25(7), 410–418.

Csilléry, K., François, O., & Blum, M. G. B. (2012). Abc: An R package for approximate Bayesian computation (ABC). Methods in Ecology and Evolution / British Ecological Society, 3(3), 475–479. Retrieved from arXiv.

Dal Vechio, F., Prates, I., Grazziotin, F. G., Zaher, H., & Rodrigues, M. T. (2018). Phylogeography and historical demography of the arboreal pit viper Bothrops bilineatus (Serpentes, Crotalinae) reveal multiple connections between Amazonian and Atlantic rain forests. Journal of Biogeography, 45(10), 2415–2426.

Ewing, G. B., Reiff, P. A., & Jensen, J. D. (2015). PopPlanner : visually constructing demographic models for simulation. 6(April), 1–4.

Fagundes, N. J. R., Ray, N., Beaumont, M., Neuenschwander, S., Salzano, F. M., Bonatto, S. L., & Excoffier, L. (2007). Statistical evaluation of alternative models of human evolution. Proceedings of the National Academy of Sciences of the United States of America, 104(45), 17614–17619.

Gehara, M., Garda, A. A., Werneck, F. P., Oliveira, E. F., da Fonseca, E. M., Camurugi, F., … Burbrink, F. T. (2017). Estimating synchronous demographic changes across populations using hABC and its application for a herpetological community from northeastern Brazil. Molecular Ecology, 26(May), 4756–4771.

Gill, M. S., Lemey, P., Faria, N. R., Rambaut, A., Shapiro, B., & Suchard, M. A. (2013). Improving bayesian population dynamics inference: A coalescent-based model for multiple loci. Molecular Biology and Evolution, 30(3), 713–724.

Gloyd, H. K., & Conant, R. (1990). Snakes of the Agkistrodon complex: a monographic review. Contributions to Herpetology, 1–614. Oxford, Ohio: Society for the Study of Amphibians and Reptiles.

Gow, J. L., Peichel, C. L., & Taylor, E. B. (2007). Ecological selection against hybrids in natural populations of sympatric threespine sticklebacks. Journal of Evolutionary Biology, 20(6), 2173–2180.

Gronau, I., Hubisz, M. J., Gulko, B., Danko, C. G., & Siepel, A. (2011). Bayesian inference of ancient human demography from individual genome sequences. Nature Genetics, 43(10), 1031–1034.

Guiher, T. J., & Burbrink, F. T. (2008). Demographic and phylogeographic histories of two venomous North American snakes of the genus Agkistrodon. Molecular Phylogenetics and Evolution, 48(2), 543–553.

Hey, J. (2010). Isolation with migration models for more than two populations. Molecular Biology and Evolution, 27(4), 905–920.

Hine, A. C. (2013). Geologic history of Florida: major events that formed the Sunshine State. Gainesville, USA: University Press of Florida.

Hudson, R. (2002). Ms a program for generating samples under neutral models. Bioinformatics, (2002), 337–338.

Kingman, J. F. C. (1982). The coalescent. Stochastic Processes and Their Applications, Vol. 13, pp. 235–248. doi: 10.1016/0304-4149(82)90011-4

Krysko, K. L., Nuñez, L. P., Lippi, C. A., Smith, D. J., & Granatosky, M. C. (2016). Molecular Phylogenetics and Evolution Pliocene – Pleistocene lineage diversifications in the Eastern Indigo Snake (Drymarchon couperi) in the Southeastern United States q. Molecular Phylogenetics and Evolution, 98, 111–122.

Kuhn, M. (2008). Building Predictive Models in R Using the caret Package. Journal of Statistical Software, 28(5), 1–26. Retrieved from arXiv.

Marshall, S. J., James, T. S., & Clarke, G. K. C. (2002). North American Ice Sheet reconstructions at the last glacial maximum. Quaternary Science Reviews, 21(1-3), 175–192.

Mayr, E. (1942). Systematics and the Origin of Species, from the Viewpoint of a Zoologist. Harvard University Press.

Mckelvy, A. D., & Burbrink, F. T. (2017). Molecular Phylogenetics and Evolution Ecological divergence in the yellow-bellied kingsnake (Lampropeltis calligaster) at two North American biodiversity hotspots. Molecular Phylogenetics and Evolution, 106, 61–72.

Nielsen, R., & Beaumont, M. A. (2009). Statistical inferences in phylogeography. Molecular Ecology, Vol. 18, pp. 1034–1047. doi: 10.1111/j.1365-294x.2008.04059.x

Nomura, F., Rossa-Feres, D. C., & Langeani, F. (2009). Burrowing behavior of Dermatonotus muelleri (Anura, Microhylidae) with reference to the origin of the burrowing behavior of Anura. Journal of Ethology, 27(1), 195–201.

Oliveira, E. F., Gehara, M., São-Pedro, V. A., Costa, G. C., Burbrink, F. T., Colli, G. R., … Garda, A. A. (2018). Phylogeography of Muller’s termite frog suggests the vicariant role of the Central Brazilian Plateau. Journal of Biogeography, 45(11), 2508–2519.

Pavlidis, P., & Laurent, S. (n.d.). msABC : A modification of Hudson ‘ s ms to facilitate multilocus ABC analysis User ‘ s Guide & Manual Table of Contents. doi: 10.1111/j.1755-0998.2010.02832.x

Pavlidis, P., Laurent, S., & Stephan, W. (2010). MsABC: A modification of Hudson’s ms to facilitate multi-locus ABC analysis. Molecular Ecology Resources, 10(4), 723–727.

Pfeifer, B., Wittelsburger, U., Ramos-Onsins, S. E., & Lercher, M. J. (2014). PopGenome: An efficient swiss army knife for population genomic analyses in R. Molecular Biology and Evolution, 31(7), 1929–1936.

Prates, I., Rivera, D., Rodrigues, M. T., & Carnaval, A. C. (2016). A mid-Pleistocene rainforest corridor enabled synchronous invasions of the Atlantic Forest by Amazonian anole lizards. Molecular Ecology, 25(20), 5174–5186.

Schrider, D. R., & Kern, A. D. (2018). Supervised Machine Learning for Population Genetics: A New Paradigm. Trends in Genetics: TIG, xx, 1–12.

Sheehan, S., & Song, Y. S. (2016). Deep Learning for Population Genetic Inference. PLoS Computational Biology, 12(3). doi: 10.1371/journal.pcbi.1004845

Soltis, D. E., Morris, A. B., McLachlan, J. S., Manos, P. S., & Soltis, P. S. (2006). Comparative phylogeography of unglaciated eastern North America. Molecular Ecology, 15, 4261–4293.

Yang, Z., & Rannala, B. (2010). Bayesian species delimitation using multilocus sequence data. Proceedings of the National Academy of Sciences of the United States of America, 107(20), 9264–9269.

